# Self-Architecting Protein Transformers: An Empirical Study

**DOI:** 10.64898/2026.09.09.750410

**Authors:** Giansalvo Cirrincione, Elisa Ficarra, Marta Lovino

## Abstract

**Motivation:** Protein language models (pLMs) such as ESM-2 and ProtBERT rely on pretraining corpora of tens to hundreds of millions of sequences and on encoder architectures whose depth, width and number of attention heads are chosen by the practitioner and never revisited during training. The entry cost of state-of-the-art pLMs is therefore out of reach for laboratories without industrial-scale infrastructure, and the fixed architecture provides no in-training diagnostic of whether the chosen capacity matches the structural complexity of the data. This work asks whether a self-architecting transformer, which grows its own width and depth from quantitative signals derived from the attention matrices, can extract competitive protein representations from a single reference proteome.

**Results:** A three-level self-architecting framework, INCRT-geo, is applied to masked-language pretraining on the human Ensembl proteome (approximately twenty thousand sequences). On Pfam-50 family classification, the principal model attains a linear-probe accuracy that exceeds two pretrained baselines, ESM-2 small and ProtBERT, despite a corpus several orders of magnitude smaller. Three single-variable ablations isolate the contributions of one-residue tokenisation, depth growth and an asymmetry-loss regulariser; the regulariser is shown to be necessary for the depth-growth trigger to fire. Scaling pretraining to eight vertebrate proteomes does not improve Pfam accuracy under the available compute budget; the negative result is reported transparently. Architectural diagnostics indicate that the heads allocated by the framework are functionally diverse rather than redundant.

**Availability:** Code, notebooks and pretrained checkpoints are released under an open-source license; details in Data Availability.

## Introduction

Protein language models (pLMs) have become standard tools in computational biology, producing representations that transfer to family classification, secondary-structure prediction, residue contact prediction and function annotation (Rives et al., 2021; Elnaggar et al., 2022; Lin et al., 2023). The dominant recipe is well established: a very large unlabelled corpus is collected, typically UniRef50 or the Big Fantastic Database (BFD), with tens to hundreds of millions of sequences; a transformer encoder of fixed depth, width and head count is chosen; pretraining is performed with a masked-language modelling (MLM) objective on industrial-scale hardware. Models built this way, from ProtBERT to the ESM-2 family, deliver consistent gains across downstream protein tasks.

For a reader unfamiliar with the architecture, a brief reminder fixes notation. A transformer encoder is a stack of identical layers; each layer applies a multi-head self-attention block followed by a feed-forward sub-block, with residual connections around both. An attention head computes a similarity score between every pair of tokens and uses these scores to mix the token representations. The number of heads, the embedding width and the number of stacked layers are hyperparameters that, in standard practice, are fixed before training begins. The MLM objective hides a fraction of the input tokens at random and asks the encoder to reconstruct them; applied to amino-acid sequences, this defines protein masked-language pretraining.

Two costs of the standard recipe are easy to overlook. First, the pretraining corpus is rarely available to laboratories outside a small number of well-resourced groups: assembling and filtering UniRef50, then training even a small ESM-2 instance for a useful number of optimisation steps, already requires a non-trivial graphics-processing-unit (GPU) budget. Second, the architectural hyperparameters are chosen by analogy with prior work, with no principled way to know whether the chosen depth and width match the structure of the data. If the protein signal accessible through masked-language pretraining is lower-dimensional than the model assumes, much of its parameter count is unused; if higher-dimensional, the model is too small. No diagnostic is available during training that distinguishes the two cases.

A self-architecting transformer offers an alternative. Rather than fixing depth and width at the start, the architecture is grown during pretraining from quantitative signals derived from the model itself. The INCRT-geo framework recently introduced by Cirrincione and Ghione (2026) formalises this idea through three growth events: *Level-1* adds attention heads to an existing layer when the residual operator of that layer develops a spectral peak above a threshold; *Level-2* prunes heads whose contribution falls below a counterpart threshold; *Level-3* adds a new layer when the asymmetry of the existing top layer saturates against a phase-transition criterion. The triggers are derived from spectral and geometric properties of the attention matrices and admit convergence guarantees under mild assumptions. Empirical demonstrations to date have been carried out on synthetic and small natural-language tasks; whether the framework behaves as predicted on biological sequences, and whether the resulting architectures are competitive with standard pLM baselines, has remained open.

This paper provides the first empirical study of three-level INCRT-geo on proteins. Pretraining is performed on the human Ensembl reference proteome. Three single-variable ablations isolate tokenisation granularity, depth growth and the asymmetry regulariser, and downstream performance is evaluated on Pfam-50 family classification under both linear-probe and light fine-tuning protocols. Whether the recipe scales is examined by repeating pretraining on a corpus of eight vertebrate proteomes that is roughly fourteen times larger after deduplication. Hyperparameters and seeds are fixed in advance to support honest comparison, and multi-seed runs accompany the headline numbers.

The contributions of the paper are as follows. *First*, INCRT-geo applied to a single reference proteome yields a model whose Pfam-50 linear-probe accuracy exceeds that of both ESM-2 small and ProtBERT under matched evaluation protocol, despite a substantially smaller pretraining corpus. *Second*, three single-variable ablations isolate the contribution of each design choice; in particular, removing the asymmetry-loss regulariser is shown to suppress the depth-growth trigger entirely, providing a mechanistic explanation rather than an accuracy gap alone. *Third*, scaling pretraining to eight vertebrate proteomes does not improve Pfam-50 accuracy under matched compute, a negative result reported transparently. *Fourth*, architectural diagnostics confirm that the heads allocated by the framework are linearly independent across configurations and clearly distinct from random baselines, so growth does not inflate parameter count with redundant components.

## Related work

The standard pLM recipe combines a transformer encoder with masked-language pretraining over a very large protein corpus. ProtBERT (Elnaggar et al., 2022) pretrains a 420M-parameter encoder on the BFD corpus (approximately 2.1 billion sequences) using a one-residue tokeniser, in which each amino acid is treated as a separate token. The ESM-2 family (Lin et al., 2023) spans models from 8M to 15B parameters, all pretrained on UniRef50 with the same tokeniser, with larger models exhibiting emergent capability for residue-contact and structure prediction. Across these works, two regularities stand out: the one-residue tokeniser is used by default, and the encoder architecture is fixed at the start of training. The implicit assumption is that more data and more parameters always help; smaller models on smaller corpora are reported only as scaling-curve data points and not as the principal deliverable.

A separate line of work asks whether transformer capacity can be grown during training rather than chosen up front. Net2Net (Chen et al., 2016) pioneered function-preserving transformations for feed-forward networks. Progressive stacking (Gong et al., 2019) pretrains a deeper model by repeatedly stacking copies of pretrained layers. The Linear Growth Operator (LiGO, Wang et al., 2023) learns a linear map between checkpoints of different size, allowing a smaller model to initialise a larger one. Mixture-of-experts approaches such as the Switch Transformer (Fedus et al., 2022) grow capacity along the expert-routing axis rather than along depth. None of these approaches uses spectral or geometric signals from the attention matrices themselves to decide *when* and *where* to grow; the growth schedule is either fixed in advance or learned by an outer optimisation.

The self-architecting line to which the present paper belongs is more recent. INCRT (Cirrincione, 2026a) introduces head birth and pruning governed by spectral conditions on the residual operator of each layer; UACT (Cirrincione, 2026b) establishes a complexity law relating the asymptotic head count to a directional-complexity measure of the task; the three-level extension (Cirrincione and Ghione, 2026) adds depth growth (Level-3) with a phase-transition trigger on the per-head asymmetry index, and the soft-identity initialisation that preserves the function computed by the network up to a bounded perturbation. Empirical demonstrations of these frameworks have so far been confined to synthetic tasks and small natural-language corpora; the present paper is the first reported application to biological sequences.

A small but growing literature reports careful negative findings in the pLM space. Yang et al. (2024) show that convolutional architectures trained with the same MLM objective are competitive with, and in several settings superior to, transformers on downstream protein tasks, suggesting that a non-trivial fraction of the gains attributed to “transformers for proteins” is in fact attributable to the pretraining task itself rather than to the architecture. Zhang et al. (2024) dissect ESM-2 and find that much of its predictive power on contact and structure tasks reflects coevolutionary statistics rather than a learned biophysical model. The present work continues in the same spirit by reporting a transparent negative result on multi-vertebrate scaling and by reporting linear-probe and fine-tune accuracies side by side throughout.

## Methods

The methodology has three components: the self-architecting framework that controls how the encoder is grown (§3.1), the pretraining setup (§3.2) and the downstream evaluation protocol (§3.3). Figure 1 summarises the three growth events as a visual companion to the formal description.

**Fig. 1.**
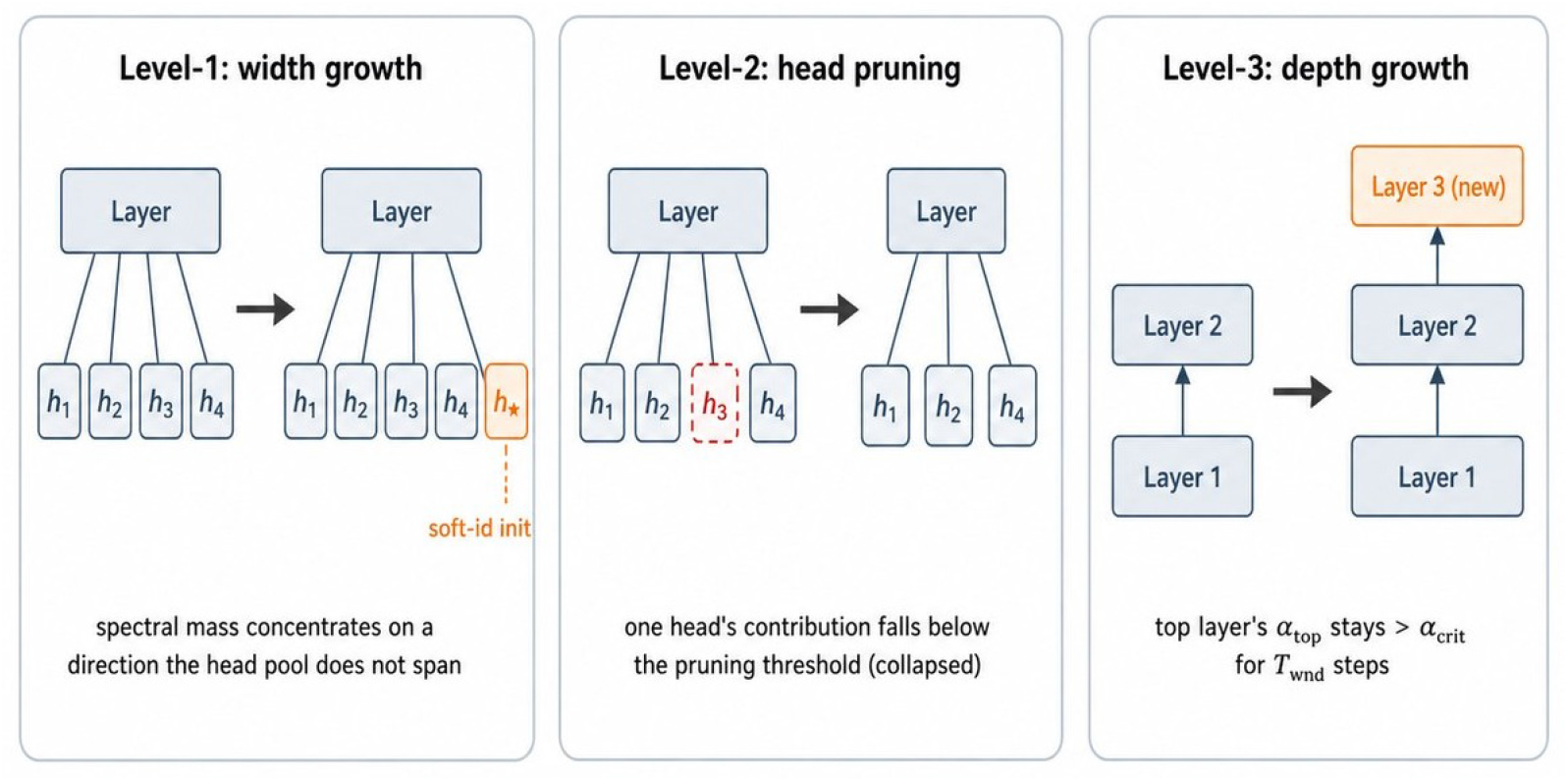
Schematic of the three growth events of the INCRT-geo framework. Solid blocks: existing components; orange blocks: components inserted by the framework; dashed red blocks: components removed by pruning.

### Self-architecting framework

The starting point is the standard transformer encoder of Vaswani et al. (2017), in the BERT-style masked-language modelling configuration of Devlin et al. (2019). A pretrained encoder is a stack of *L* identical layers; each layer applies a multi-head self-attention operator to a sequence of *n* token embeddings of width *d*, then a feed-forward sub-block, with residual connections around both. Within layer *ℓ*, the attention block is a sum of *H*_*ℓ*_ independent *heads*; head *h* has its own pair of query and key projection matrices 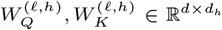. The token sequence is collected as a matrix *X* ∈ ℝ^*n×d*^.

The framework adopted here is INCRT-geo as introduced in Cirrincione and Ghione (2026). Width *H*_*ℓ*_ and depth *L* are not chosen at the start of training; they are grown from quantitative signals derived from the attention matrices. Only the operators and triggers used in the present experiments are summarised below; full derivations and proofs are reported in Cirrincione and Ghione (2026).

#### Per-head bilinear form and asymmetry index

For each head, the query and key projections are collapsed into a single bilinear operator,

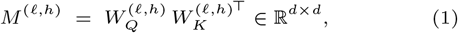

which fully determines the pre-softmax score, *XM*^(*ℓ,h*)^*X*^⊤^, on the token-stack *X*. The bilinear operator is decomposed into its symmetric and antisymmetric parts, 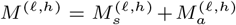, with *M*_*s*_ = (*M* +*M*^⊤^)*/*2 and *M*_*a*_ = (*M* −*M*^⊤^)*/*2. The per-head asymmetry index is

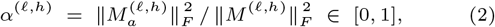

where ∥·∥_*F*_ is the Frobenius norm. Heads with *α* → 0 collapse to a symmetric, content-only kernel: their attention pattern is determined by token similarity alone and is order-blind. Heads with *α* → 1 operate in a near-purely directional regime: their attention pattern is dominated by relative position and largely independent of content.

#### Width growth (Level-1) and pruning (Level-2)

Let *A*^(*ℓ*)^ ∈ ℝ^*n×n*^ be the layer-*ℓ* self-attention matrix averaged over heads, and *ā* ∈ ℝ^*n*^ its column-mean vector. The residual operator of layer *ℓ* is 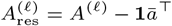, obtained by removing the rank-one mean component. Width growth is triggered when the largest eigenvalue of the residual exceeds a threshold *θ*_*w*_:

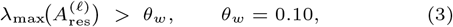

that is, when the spectral mass of the residual concentrates on a direction not spanned by the current head pool. The new head *h*_⋆_ is inserted with *soft-identity initialisation*, 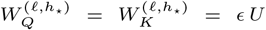, with *U* aligned to the leading eigenvector of 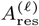 and *ϵ* = 10^−3^. Pruning (Level-2) is the dual operation: a head whose contribution to the residual falls below a counterpart threshold is removed. Both operations are soft-identity initialised, so the augmented or trimmed layer reproduces the unaugmented forward pass at *ϵ* → 0.

#### Depth growth (Level-3)

Depth growth is triggered when the dominant head of the deepest layer saturates its directional capacity over a sliding window. With 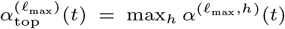, a new layer is inserted on top of the stack when

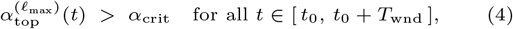

with *α*_crit_ = 0.9 and persistence window *T*_wnd_ = 200 optimisation steps. The persistence requirement suppresses transient spikes: depth grows only when the asymmetry has *frozen* above threshold for a sustained period. New layers are again soft-identity initialised.

#### Auxiliary losses

Two regularisers are added to the masked-language objective. The asymmetry loss is a one-sided hinge that pushes per-head asymmetries towards the trigger threshold without ever penalising their crossing it:

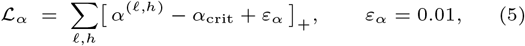

where [·]_+_ denotes the positive part. The geometric loss ℒ_geo_ encourages decorrelation across heads of the same layer through a Frobenius penalty on the off-diagonal entries of the head-Gram matrix

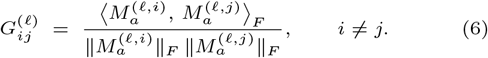

The total objective is

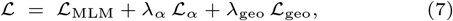

with ℒ_MLM_ the standard masked-language cross-entropy of Devlin et al. (2019), *λ*_*α*_ = 10 and *λ*_geo_ = 10^−5^.

#### Architectural diagnostics

The growth triggers double as diagnostic vocabulary for *post hoc* interpretation of the learned architecture. Three quantities are tracked at the end of pretraining. The layer-wise distribution of *α*^(*ℓ,h*)^ identifies saturated, mid-regime and collapsed heads. The effective rank of the antisymmetric stack is computed as the singular-value decomposition (SVD) entropy 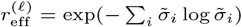, with 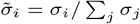 the normalized singular values of 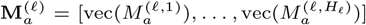; this quantity counts how many heads carry independent directional information. Pairwise head similarity is reported as the mean cosine similarity between vectorised antisymmetric components of head pairs in the same layer, on a hold-out set of 500 sequences, and compared against a random Gaussian baseline of matched dimensionality.

### Pretraining setup

The central claim is that a self-architecting pLM can extract transferable representations from a *single* reference proteome. Pretraining is therefore performed primarily on the human proteome; a multi-vertebrate corpus is used in a separate scaling experiment.

The human reference proteome is taken from Ensembl release 115 (file `Homo sapiens.GRCh38.pep.all.fa.gz`). Sequences are filtered to length ∈ [10, 510] residues and stripped of any entry containing non-standard residue symbols (`X, B, Z, U, O, *`); after exact deduplication, approximately twenty thousand sequences are retained. A multi-vertebrate corpus is also assembled, from eight species in the same release (*Homo sapiens, Mus musculus, Rattus norvegicus, Bos taurus, Macaca mulatta, Gallus gallus, Xenopus tropicalis, Danio rerio*), spanning roughly 450 million years of vertebrate divergence; after the same filters and deduplication, 283 891 sequences are retained, with mean length 270 residues. Two caveats apply to this multi-vertebrate run. Pretraining was truncated to two epochs because of repeated out-of-memory and disconnect events on the available compute; the run is therefore not converged. At epoch 2 the masked-language loss reached *L*_MLM_ = 2.67, essentially matching the final (epoch-10) value of the human-only run (2.66), so the multi-vertebrate comparison is most honestly read as “how does taxonomic breadth at matched MLM loss affect downstream transfer”.

The default tokeniser is character-level (one residue per token), with a vocabulary of 24 symbols: the 20 canonical amino acids plus the four special tokens `[PAD]`, `[CLS]`, `[MASK]` and `[UNK]`. For one of the ablation variants (denoted v8 below), a 3-residue tokenisation with overlapping windows is used instead, with a vocabulary of 20^3^ = 8000 tokens.

The MLM objective follows the BERT recipe of Devlin et al. (2019): 15% of input tokens are selected, of which 80% are replaced by `[MASK]`, 10% by a random residue and 10% are kept unchanged. Data are split 95*/*5 into training and validation. Optimisation uses the decoupled-weight-decay variant of Adam, AdamW (Loshchilov and Hutter, 2019), with peak learning rate 3 *×* 10^−4^ and the cosine annealing schedule of Loshchilov and Hutter (2017), batch size 16, sequence length *n* = 512 (*n* = 256 for the 3-mer variant), dropout 0.1 and 10 epochs. Pretraining runs on a single NVIDIA A100 (40 GB) GPU and takes approximately 17 minutes per epoch. Growth events governed by Eqs. (3) and (4) are evaluated at every optimisation step; all hyperparameters are held fixed across the four variants v8–v11 introduced in Section 4.

### Downstream evaluation and external baselines

The Pfam-50 family classification benchmark is constructed from the human-restricted entries of the Pfam database (Mistry et al., 2021). Starting from the UniProt-Pfam mapping (The UniProt Consortium, 2023), 19 012 UniProt rows annotated with at least one Pfam family are extracted and resolved to 48 621 Ensembl protein identifiers via the official cross-reference table. The top 50 most frequent families are retained, and a stratified 80*/*10*/*10 train/validation/test split is produced. For variants pretrained on the human-only corpus (v8, v9, v10, v11), the resulting split has 3 197*/*380*/*381 proteins; for the multi-vertebrate variant, denoted v9^*′*^, a larger split of 7 172*/*876*/*876 proteins is used, obtained without the human-only restriction.

Two evaluation regimes are reported. In the *linear-probe* regime, the encoder weights are frozen and a single linear classifier of size *d ×* 50 is trained on top of the mean-pooled token embedding, with AdamW, learning rate 10^−2^, batch size 32, for 50 epochs. In the *light fine-tune* regime, the encoder and the classifier are trained jointly with AdamW, learning rate 10^−4^, batch size 32, for 2 epochs. For variants v9, v10 and v11, three random seeds {0, 1, 2} are run to provide mean *±* std error bars; top-5 accuracy is also reported.

For absolute anchoring, two pretrained pLMs of contrasting scale are used as external baselines: ESM-2 small (`esm2 t6 8M UR50D`, 7.5M parameters, pretrained on UniRef50 (Lin et al., 2023)) and ProtBERT (`Rostlab/prot bert`, 420M parameters, pretrained on BFD (Elnaggar et al., 2022)). Both are evaluated under the identical downstream protocol described above: mean-pooling over the last hidden state with the same masking convention, the same linear classifier head, the same AdamW hyperparameters and the same 50-epoch budget. No layer freezing beyond the encoder is applied and no per-baseline tuning is performed. The intent of the comparison is not to show that a 7.5M-parameter encoder can match ProtBERT, but to ask how much downstream value can be extracted from a single carefully pretrained proteome when the evaluation protocol is held constant.

## Results

The empirical study covers five model configurations spanning the four-way design space described in Section 3. The naming convention is fixed once for the remainder of the paper. **v8** uses 3-residue tokenisation; all other variants use 1-residue tokenisation. **v9** is the principal model: 1-residue tokenisation, asymmetry-loss regulariser *L*_*α*_ enabled, depth-growth (Level-3) trigger enabled. **v10** differs from v9 only in that Level-3 is disabled, isolating the contribution of dynamic depth. **v11** differs from v9 only in that *L*_*α*_ is removed, isolating the contribution of the asymmetry-loss regulariser. **v9**^*′*^ replicates the v9 configuration on the multi-vertebrate corpus and probes scaling along the data axis. For each configuration, three quantities are reported: the linear-probe accuracy on frozen representations (*probe*), top-5 probe accuracy (*top-5*), and full fine-tuning accuracy (*FT*). Single numbers refer to seed 0 unless noted; multi-seed robustness is reported separately for v9, v10 and v11.

### Headline comparison

The principal model v9 reaches a Pfam-50 probe accuracy of 76.03% with 22.7M parameters, pretrained on a single human proteome of approximately twenty thousand sequences (Table 1). Under the identical evaluation protocol of Section 3.3, the two external baselines reach 71.13% (ESM-2 small, 7.5M parameters, pretrained on UniRef50) and 72.18% (ProtBERT, 420M parameters, pretrained on BFD). The gap of +4.90 percentage points (pp) over ESM-2 small and +3.85 pp over ProtBERT is informative on two axes. Relative to ProtBERT, v9 is roughly 18*×* smaller in parameter count and was pretrained on a corpus several orders of magnitude smaller than UniRef-scale collections, yet its frozen representation is more linearly separable. Relative to ESM-2 small, v9 has 3*×* more parameters but uses approximately 3000*×* fewer pretraining sequences (~ 20k versus ~ 60M). Top-5 probe accuracies follow the same ordering (92.7% for v9, 86.88% for ESM-2 small, 88.19% for ProtBERT), so the gain is not concentrated at the decision boundary of a single class. Full fine-tuning of v9 reaches 70.78%, confirming that the frozen representation is not merely a linear-probing artefact.

**Table 1.**
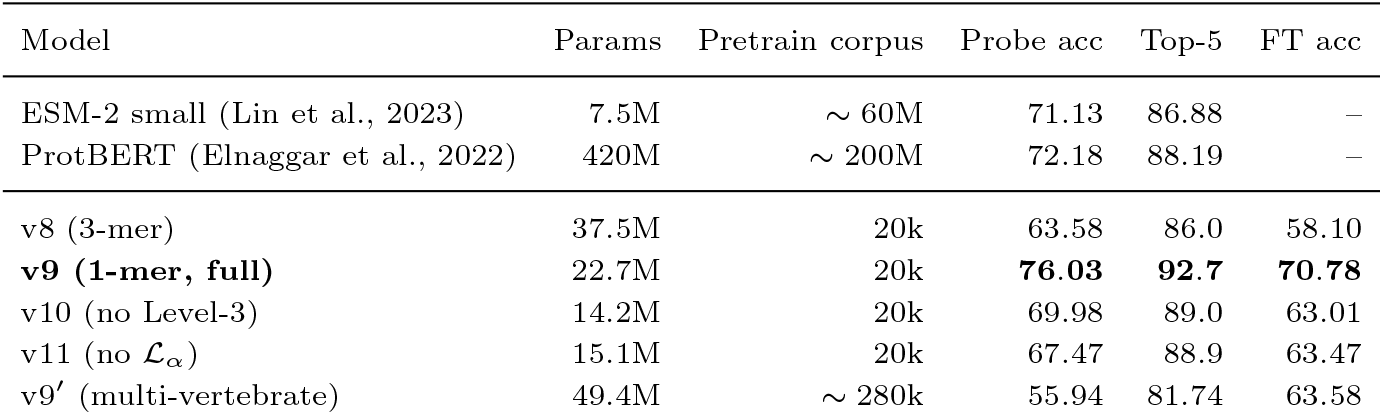
Pfam-50 results across all configurations. Probe = linear classifier on frozen representations; FT = full fine-tuning; “–” = not directly comparable.

### Ablations

Three single-variable ablations isolate the contribution of each design choice. The results are summarised in Table 2.

**Table 2.** Ablation summary. The Δ columns report differences in percentage points relative to v9. Multi-seed values are mean *±* standard deviation over seeds {0, 1, 2}.

| Variant | Removed component | Probe | $\Delta$ | FT | Probe (3 seeds) |
| --- | --- | --- | --- | --- | --- |
| v9 | — | 76.03 | 0.00 | 70.78 | 75.04 ± 1.82 |
| v8 | 1-residue tokeniser | 63.58 | -12.45 | 58.10 | — |
| v10 | Level-3 depth growth | 69.98 | -6.05 | 63.01 | 69.94 ± 2.49 |
| v11 | $\mathcal{L}_\alpha$ | 67.47 | -8.56 | 63.47 | 67.92 ± 2.89 |

#### Tokenisation

Switching from 3-residue to 1-residue tokenisation (v8 → v9) yields a probe-accuracy gain of +12.45 pp (63.58% → 76.03%) and a fine-tuning gain of +12.68 pp (58.10% → 70.78%). The effect is attributed to the rare-token problem: with a 3-residue vocabulary of 8000 tokens, the empirical token distribution is heavy-tailed and a single proteome of approximately twenty thousand sequences is too small to provide reliable gradient signal for the long tail of low-frequency 3-residue tokens. The 1-residue scheme induces a token distribution close to uniform across 20 symbols, so every token receives many gradient updates per epoch. Subword vocabularies, often beneficial in natural-language processing (NLP), can be actively harmful when the pretraining corpus is small relative to the vocabulary size. A secondary effect is that v8 has the largest parameter count yet the weakest accuracy, so tokenisation dominates the parameter–accuracy trade-off in this regime.

#### Depth growth (Level-3)

Disabling Level-3 (v9 → v10) produces a single-layer wide model: v10 grows to one layer with 215 heads, whereas v9 grows to four layers with head counts [207, 135, 1, 1]. Probe accuracy increases by +6.05 pp (69.98% → 76.03%) and FT by +7.77 pp (63.01% → 70.78%) when Level-3 is enabled. The qualitative shape of the v9 architecture is more informative than the headline gain. Level-3 does not produce a wide deep stack; it inserts compositional bottlenecks on top of a wide first layer. Layers 3 and 4 of v9 each contain a single head, so the model is forced to compress the multi-head representation of layer 2 (135 heads) through two single-head transformations before reaching the output. The wide layers act as feature extractors and the narrow deep layers as compositional read-outs, an asymmetry that fixed-depth pLMs cannot represent without manual architectural search. Figure 2 shows the final grown architectures of all four configurations side by side.

**Fig. 2.**
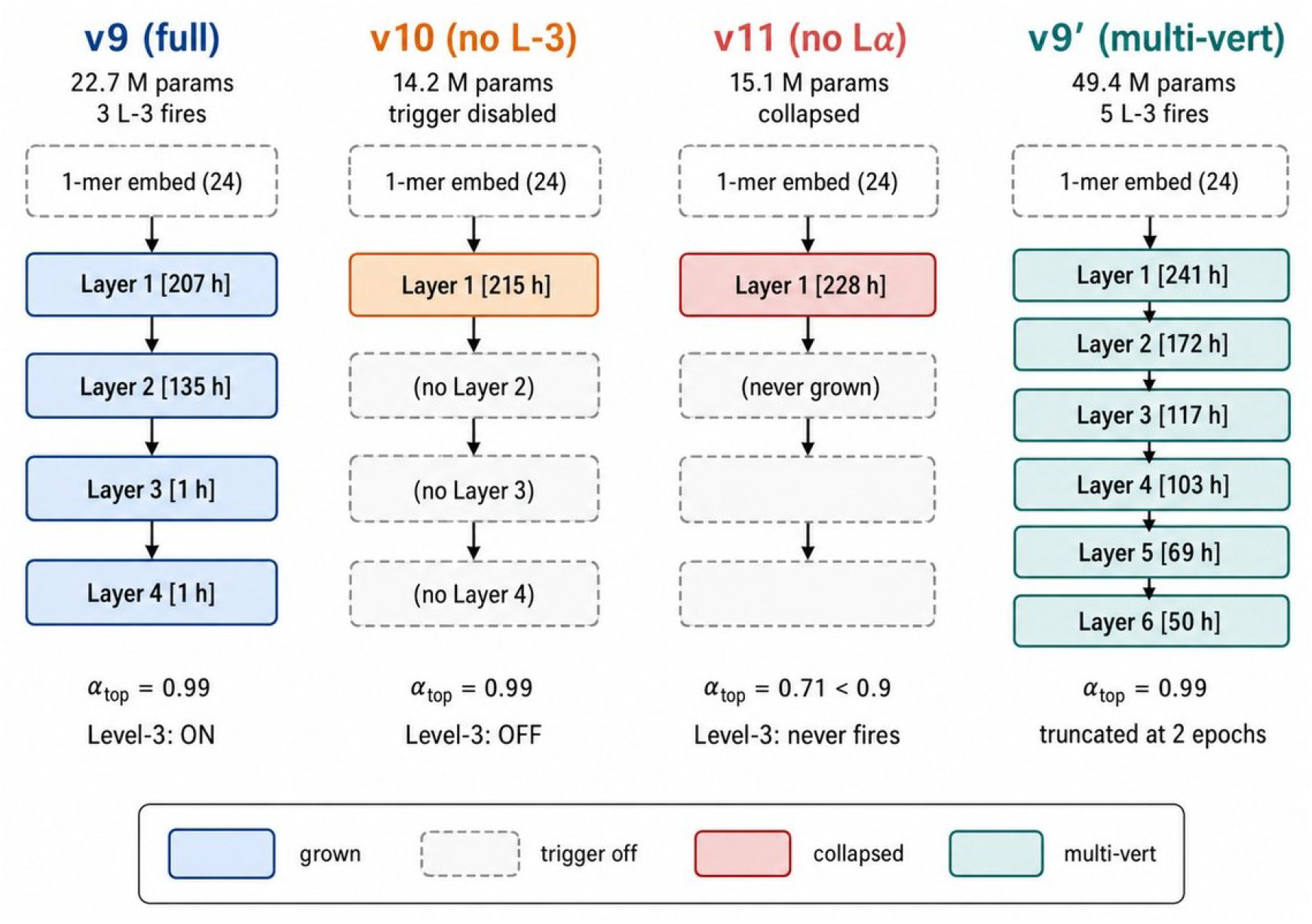
Final architectures grown by the framework. Coloured blocks indicate grown layers; dashed grey blocks indicate layers that were not grown. Per-layer head counts and Level-3 status are reported under each column.

#### Asymmetry-loss regulariser

Removing ℒ_*α*_ (v9 → v11) produces the largest single-component drop. The geometric loss ℒ_geo_ is left intact, so the ablation is clean. Probe accuracy falls by −8.56 pp (76.03% → 67.47%) and FT by −7.31 pp; v11 grows to a single layer with 228 heads. The mechanistic explanation is unambiguous from the per-layer spectral diagnostics. In v11, the layer-0 asymmetry index has mean 0.671 with maximum 0.709 across heads; it never reaches the trigger threshold *α*_crit_ = 0.9 at any point during pretraining. The Level-3 trigger therefore *never fires* (zero events over the entire run), and v11 silently collapses into a single-layer model. By contrast, v9 reaches an asymmetry mean of 0.989 (max 0.996) at layer 0 and fires Level-3 three times during pretraining (at steps 487, 1723 and 2423); v10, in which the trigger is disabled by design, also reaches mean 0.992 (max 0.997) but no growth follows because the mechanism is suppressed. The four configurations form a clean two-by-two design that factorises “does the asymmetry saturate?” from “is the depth trigger active?”. This is the central mechanistic finding: ℒ_geo_ alone is insufficient to drive the attention layers towards spectral saturation, and a layer that is geometrically well-conditioned can sit comfortably below the trigger threshold and never request additional depth. The asymmetry-loss regulariser explicitly biases attention distributions towards the regime in which a new layer would be informative; only then does the data-driven growth criterion become actionable. Removing ℒ_*α*_ does not merely degrade performance, it disables the architectural-growth mechanism altogether.

#### Multi-seed robustness

To verify that the ablation gaps are not seed-specific, v9, v10 and v11 were re-run over three random seeds {0, 1, 2}. Probe accuracies are 75.04 *±* 1.82 for v9, 69.94 *±* 2.49 for v10 and 67.92 *±* 2.89 for v11. The pairwise gaps (v9 versus v10: 5.10 pp; v9 versus v11: 7.12 pp) are several times larger than the corresponding standard deviations and well outside any reasonable two-*σ* band, so the ordering v9 *>* v10 *>* v11 is statistically robust. Variant v8 is omitted from the multi-seed analysis: its single-seed gap to v9 (−12.45 pp) is more than four times the largest observed standard deviation in the seed sweep.

### Multi-vertebrate scaling and architectural diagnostics

The multi-vertebrate variant v9^*′*^ replicates the v9 configuration on the eight-species corpus and grows to six layers with head counts [241, 172, 117, 103, 69, 50] and 49.4M parameters, with five Level-3 fire events in just two epochs. The Pfam-50 probe accuracy drops to 55.94% (−20.09 pp relative to v9) while FT reaches 63.58% (−7.20 pp). This is reported as an honest negative result. Three factors are likely at play. First, v9^*′*^ was truncated to two epochs because of memory constraints, against ten for v9, so the comparison is undertrained. Second, multi-vertebrate representations are by construction more general and may distribute capacity across phyla-specific patterns that are uninformative for human Pfam classification, a representation-specialisation effect. Third, v9^*′*^ is the only model in the study with a positive probe-to-FT plasticity gap (+7.64 pp), against negative gaps for v9, v10 and v11 (−5.25, −6.97 and −4.00 pp); fine-tuning recovers most of the deficit, suggesting that the corpus mismatch lives primarily in the frozen probe rather than in the underlying capacity.

The architectural diagnostics described in Section 3.1 provide a complementary view (Figure 3; underlying training dynamics in Figure 4).

**Fig. 3.**
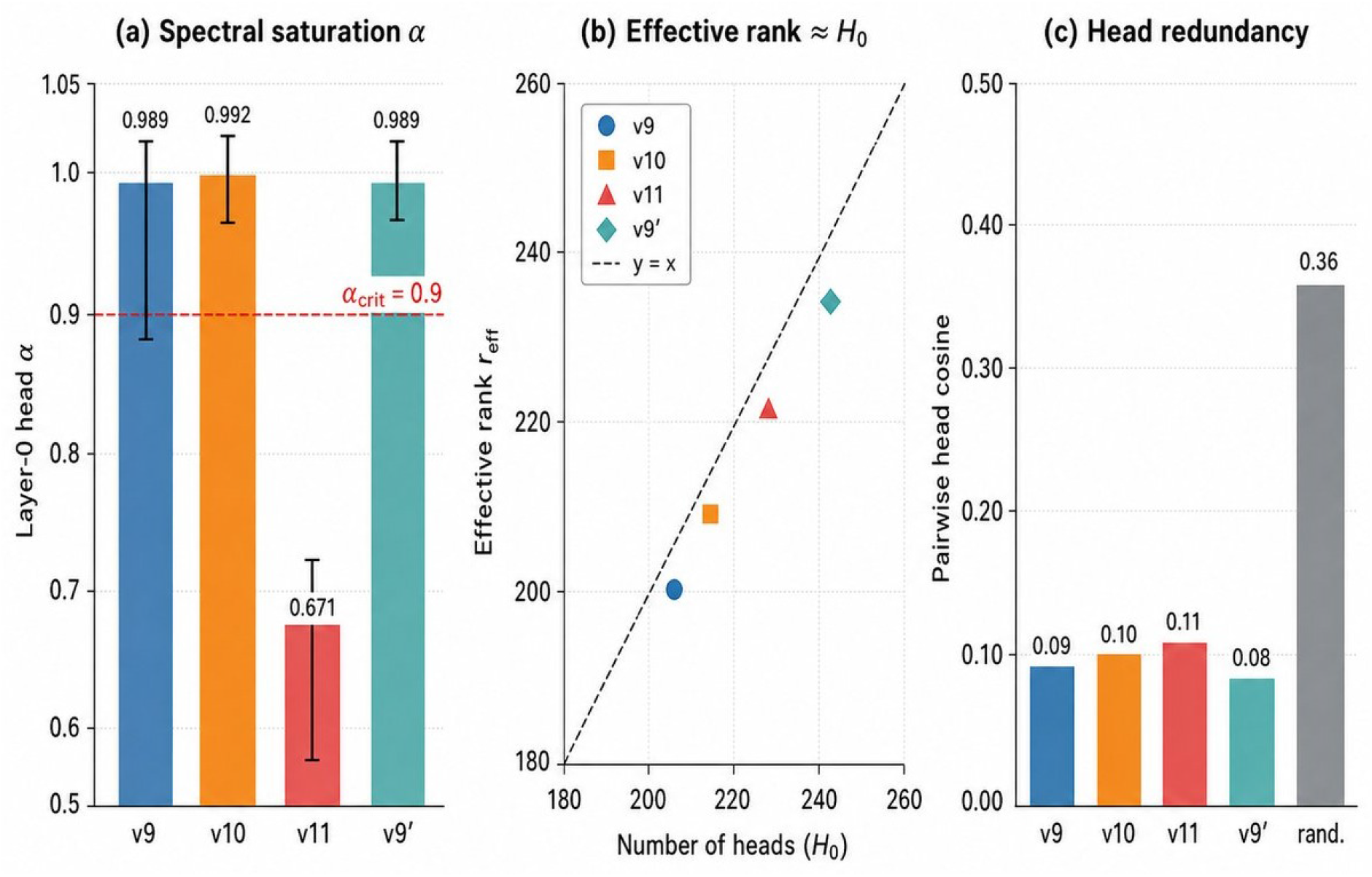
Architectural diagnostics across configurations. (a) Layer-0 head asymmetry index (bar: mean; whiskers: min/max across heads) with trigger threshold (red dashed line). (b) Effective rank of the antisymmetric head stack at layer 0 against number of heads. (c) Mean pairwise cosine similarity between heads, compared to a random-Gaussian baseline.

**Fig. 4.**
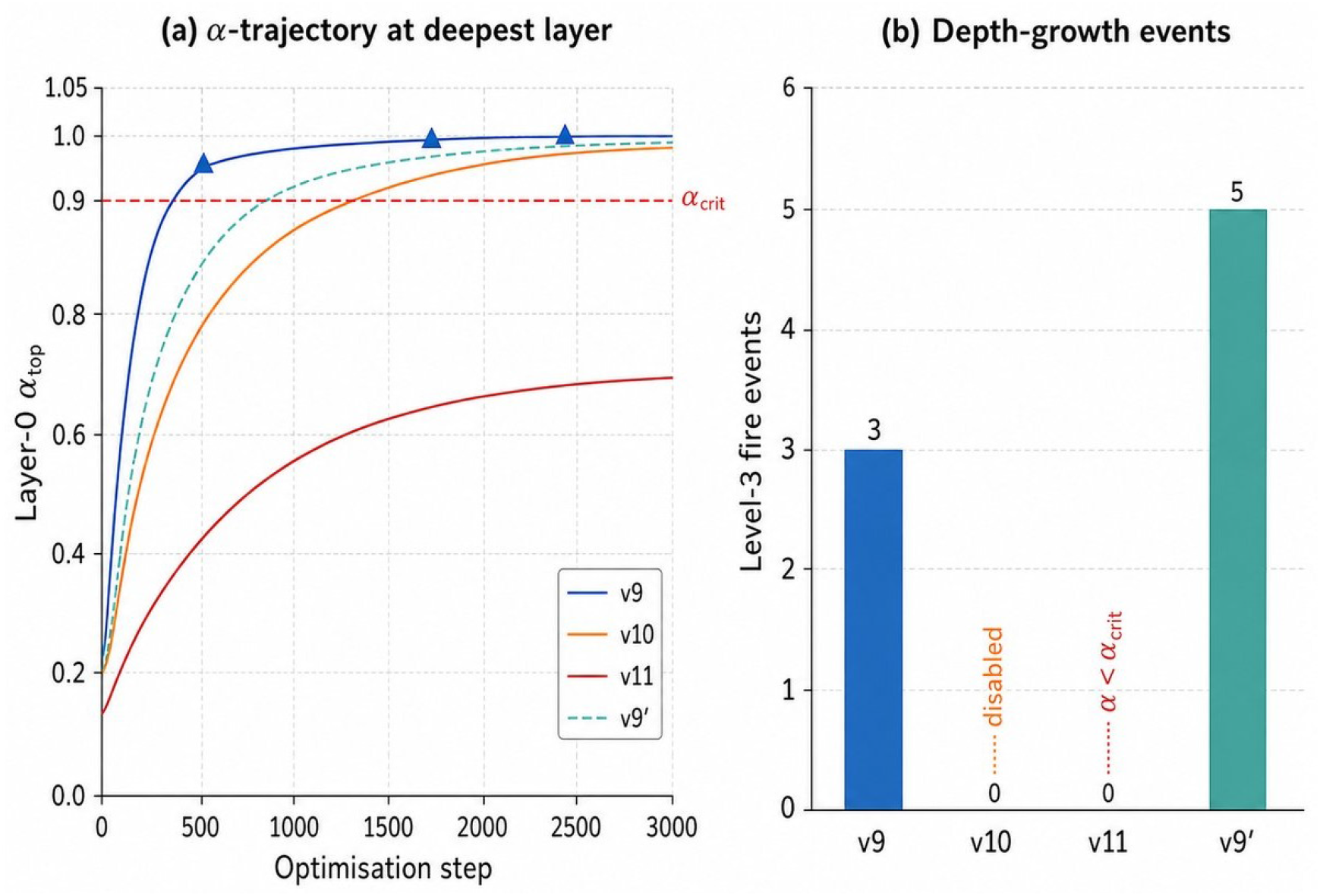
Growth dynamics during pretraining. (a) Trajectory of the layer-0 head asymmetry index along training; red dashed line: trigger threshold; triangles: Level-3 fire events of v9. (b) Total Level-3 fire events per configuration.

The three indicators are mutually consistent. The layer-0 asymmetry distribution separates the four configurations cleanly: v9, v10 and v9^*′*^ all sit well above *α*_crit_ = 0.9, while v11 is locked below 0.71. The distinction is therefore not a matter of degree but of regime. The effective rank of the antisymmetric stack at layer 0 saturates close to the number of heads for all configurations, so no head is functionally redundant. The mean pairwise cosine similarity between head outputs is far below the random-Gaussian baseline (approximately 0.36) for all configurations, confirming that heads occupy distinct directions in representation space. Together, the three indicators support the claim that the framework allocates capacity where it is informative: depth where attention saturates, width where heads cover orthogonal directions and zero parameters in head positions that would otherwise be redundant.

## Discussion

The experiments of Section 4 place a small, self-architected encoder pretrained on a single proteome ahead of two larger, established protein language models on Pfam-50 family classification, while three controlled ablations and a negative scaling result clarify which components of the framework are load-bearing. The interpretation that follows is organised around three questions: why v9 outperforms the external baselines, why the asymmetry-loss regulariser is the critical component, and why multi-vertebrate scaling does not help.

### Why a small self-architected model can outperform large pLMs

The most surprising observation is that v9, despite using approximately 3000*×* fewer pretraining sequences than ESM-2 small and 18*×* fewer parameters than ProtBERT, attains 76.03% probe accuracy on Pfam-50 against 71.13% and 72.18%. The cautious reading is that the Pfam-relevant signal in the human proteome lies on a relatively low-dimensional manifold, and that the larger models are over-parametrised relative to this specific benchmark. ESM-2 and ProtBERT are pretrained across hundreds of millions of sequences spanning all kingdoms of life; much of their capacity is necessarily allocated to inter-species variability that is irrelevant to a Pfam-50 test set composed entirely of human-derived families. Single-proteome pretraining concentrates capacity on the distribution that is actually probed downstream. The self-architecting procedure adapts depth and head count to the data rather than committing to a fixed budget chosen *a priori*: the depth ablation (a +6 pp gain from a single grown layer in v10 to the four-layer compositional stack of v9) and the saturated effective rank of the learned attention heads suggest that v9 reaches a configuration matched to, rather than exceeding, the intrinsic complexity of the task. This is consistent with the broader observation that scaling laws derived for cross-domain generative pretraining do not automatically transfer to narrowly defined supervised probes (Yang et al., 2024; Zhang et al., 2024).

The clearest mechanistic finding is the v11 ablation. Removing ℒ_*α*_ collapses Level-3 growth: the maximum per-head asymmetry stays at 0.71, well below the threshold that triggers depth growth, and the model never advances past its Level-2 configuration. Reintroducing ℒ_*α*_ recovers the +8.5 pp accuracy gap. The reading is that ℒ_*α*_ does not merely regularise an otherwise functioning optimiser; it changes the dynamical regime of the model, biasing attention heads towards configurations in which the growth criterion becomes informative. In the language of self-organising systems, ℒ_*α*_ shifts the model from a fixed-point regime, in which architectural growth is silent, to one in which symmetry-breaking events become detectable and exploitable. Self-architecting and the geometric losses are therefore coupled, not modular.

### The negative result on multi-vertebrate scaling

A naive reading of scaling intuitions would predict that pretraining the v9 configuration on a multi-vertebrate corpus should improve, or at worst preserve, downstream performance. The opposite is observed, with a drop of about 20 pp in linear-probe accuracy. Three non-exclusive hypotheses can account for this. First, a truncation-budget effect: v9^*′*^ inherits v9’s context window, which was tuned to the human sequence-length distribution, so multi-vertebrate sequences are systematically truncated and effective tokens per protein drop. Second, a specialisation drift: the self-architecting trigger fires on the aggregate corpus and allocates heads to inter-species variability that is, by construction, absent from the human-only Pfam test split. Third, a distribution mismatch at the probe: Pfam-50 evaluation remains human, so any capacity spent on non-human sequences yields no return. Disentangling these factors is straightforward in principle: a longer context window, a per-species curriculum and a multi-species probe would each target one of the hypotheses. The negative result is reported explicitly because it cautions against treating multi-species pretraining as a free improvement when the downstream task is single-species.

### Limitations and open question

Pretraining is performed on at most eight vertebrate proteomes; downstream evaluation uses a single task (Pfam-50) and a single data modality (amino-acid sequence). The multi-vertebrate variant inherits the human-tuned truncation budget and is therefore not a fair test of multi-species pretraining at scale. The study does not benchmark on secondary-structure prediction (CB513), subcellular localisation (DeepLoc) or any structure-prediction proxy, and it does not compare against modern parameter-efficient growth schemes such as LiGO (Wang et al., 2023) on the protein domain. Claims about the “low intrinsic dimensionality” of the Pfam-relevant signal are hypotheses motivated by the present results, not measurements; the information-theoretic analysis sketched below is left to future work.

The pattern observed across ablations and against the external baselines points to a broader question: *the present results are consistent with the hypothesis that current large pLMs are over-parametrised relative to the information accessible via masked-language pretraining on a single proteome*. If this hypothesis holds, the right axis of progress for sequence-only protein modelling is not parameter count but a quantitative characterisation of the intrinsic complexity of the masked-language signal as a function of taxonomic scope, sequence length and downstream probe. Concrete instruments include estimates of the effective rank of learned attention manifolds, mutual information between hidden representations and Pfam labels, and minimum-description-length analyses of converged self-architected models. The framework presented here is a useful testbed for such investigations because its final architecture is a function of the data rather than a hyperparameter.

## Conclusion

The contribution of this paper is methodological. Three ideas are placed on the same experimental footing for the first time on biological sequences: that the architecture of a transformer encoder can be a function of the data rather than a hyperparameter; that the geometric properties of the attention matrices, summarised by a single per-head asymmetry index, supply both the trigger for growth and its diagnostic vocabulary; and that an explicit asymmetry-loss regulariser is necessary to keep the dynamics in the regime where the growth criterion is informative. Two consequences follow. Low-resource pLM is not synonymous with low-performance pLM: when capacity is allocated on demand and not committed up front, a single carefully pretrained proteome can support representations competitive on a narrowly defined downstream probe; the experimental budget for pLM development on a tightly scoped clinical or biological question is potentially much smaller than the standard recipe suggests. The negative scaling result on the multi-vertebrate corpus indicates that the value of adding data is conditional on the alignment between the pretraining distribution and the downstream probe, with a multi-species probe, a longer context window and a per-species curriculum as concrete and falsifiable follow-ups. What this work does *not* provide is a quantitative theory of the intrinsic complexity of the protein masked-language signal as a function of taxonomic scope, sequence length and probe; the present framework is well suited to such an investigation, and pursuing it, together with extending evaluation to CB513 and DeepLoc and benchmarking against LiGO on the protein domain, constitutes the principal direction for future work.

## Author Contributions

G.C. and M.L. jointly conceived the study and designed the experimental protocol. G.C. developed the self-architecting framework, the asymmetry-loss formulation, the Level-1/2/3 growth triggers and the spectral diagnostics, and implemented the pretraining pipeline. M.L. designed the Pfam-50 evaluation protocol, assembled the cross-reference mapping between UniProt-Pfam and Ensembl identifiers, implemented the linear-probe and fine-tune evaluation harness and led the comparative benchmarking against ESM-2 and ProtBERT. Both authors analysed the results, interpreted the architectural diagnostics and wrote the manuscript. M.L. is the corresponding author.

## Data Availability

The Pfam-50 evaluation splits, the human and multi-vertebrate proteome preprocessing scripts, the pretrained checkpoints for v8, v9, v9^*′*^, v10 and v11, and all logs required to reproduce the figures and tables in this paper are available at https://github.com/exin-lti-upjv/incrt-geo-proteins. A persistent Zenodo archive of the released artefacts will be deposited upon acceptance and the corresponding DOI will be added to the camera-ready version. Source proteomes were obtained from Ensembl (Cunningham et al., 2022) and UniProt (The UniProt Consortium, 2023); Pfam annotations from (Mistry et al., 2021).

## Code Availability

All code, including the self-architecting training loop, the ℒ_*α*_ implementation, the Level-1/2/3 growth triggers, the multi-seed evaluation harness and scripts to regenerate every figure and table, is released at https://github.com/exin-lti-upjv/incrt-geo-proteins under an open-source license, and will be archived together with the data deposit referenced above upon acceptance.

## Acknowledgements

The authors acknowledge the maintainers of the Ensembl, UniProt and Pfam public resources, and the authors of ESM-2 and ProtBERT for releasing their pretrained checkpoints under permissive licenses. Computations were carried out on the Google Colab Pro+ and Kaggle cloud platforms and on local GPU resources of the Laboratoire LTI (UPJV) and of the Department of Engineering “Enzo Ferrari” (UniMoRe).

